# Common variants in *ABCG8* and *TRAF3* genes confer risk for gallstone disease and gallbladder cancer in admixed Latinos with Mapuche Native American ancestry

**DOI:** 10.1101/265728

**Authors:** Bernabé I. Bustos, Eduardo Pérez-Palma, Stephan Buch, Lorena Azócar, Eleodoro Riveras, Giorgia D. Ugarte, Mohammad Toliat, Peter Nürnberg, Wolfgang Lieb, Andre Franke, Sebastian Hinz, Greta Burmeister, Witigo von Schönfels, Clemens Schafmayer, Henry Völzke, Uwe Völker, Georg Homuth, Marcus M. Lerch, José Luis Santos, Klaus Puschel, Claudia Bambs, Rodrigo A. Gutiérrez, Jochen Hampe, Giancarlo V. De Ferrari, Juan Francisco Miquel

## Abstract

**Background:** Latin Americans and Chilean Amerindians have the highest prevalence of cholesterol gallstone disease (GSD) and gallbladder cancer (GBC) in the world. A handful of loci have been associated with GSD in populations of predominantly European ancestry, however they only explain a small portion of the population-attributable risk of the disease.

**Methods:** We performed a genome-wide association study (GWAS) for GSD in 1,095 admixed Latinos with Mapuche Native American Ancestry, followed by a replication analysis of 10 candidate single nucleotide polymorphisms (SNPs) with suggestive genome-wide significance (P<1×10^−5^) in 1,643 individuals. Disease status was assessed by cholecystectomy or abdominal ultrasonography. Logistic regression analyses were adjusted for age, sex, BMI, Type 2 Diabetes and Amerindian ancestry. Associated variants were further examined in two large GSD European populations and in a Chilean gallbladder cancer (GBC) cohort. We determined the expression levels of a novel GSD-candidate gene in normal and GSD-tissue samples.

**Results:** We consistently replicated the *ABCG8* gene (rs11887534; P=3.24×10^−8^, OR=1.74) associated with GSD in admixed Latinos and identified a novel candidate signal within the *TRAF3* gene on chromosome 14 (rs12882491; P=1.11×10^−7^, OR=1.40). *ABCG8* and *TRAF3* variants also conferred risk to GBC. Gene expression analyses indicated that *TRAF3* levels were significantly decreased in the gallbladder (P=0.015) and the duodenal mucosa (P=0.001) of affected GSD individuals compared to healthy controls.

**Conclusions:** We confirmed *ABCG8* and identified *TRAF3* both associated with GSD and GBC in admixed Latinos. Decreased TRAF3 expression levels could enhance gallbladder inflammation as is observed in GSD and GSD-associated GBC.

## INTRODUCTION

Gallstone disease (GSD) is a complex gastrointestinal disorder defined by the development of gallstones in the gallbladder, most of the time cholesterol compounds.^1^ The presence of gallstones are common in the world population (~10-20% of presence in adults) and although their appearance can remain silent throughout life, more than 20% of the patients develop symptoms that include intense abdominal pain, jaundice, fever, nausea and vomiting, requiring medical attention and therefore establishing itself as a clinical condition.^2^ Main complications of GSD are acute cholecystitis, acute pancreatitis and bile duct obstruction,^3^ and these are generally treated by surgical removal of the gallbladder (cholecystectomy). In Chile, these procedures accounts for more than 40,000 interventions each year,^4^ with a net cost of more than US $25 million for the public healthcare system. GSD is also the main risk factor for gallbladder cancer (GBC), a disease that presents a high mortality rates in Chile (38.2 deaths per 100,000 inhabitants).^5^ In the vast majority of cases, GBC is an adenocarcinoma characterized by high lethality due to late diagnosis and the ineffectiveness of chemotherapy/radiotherapy. With the removal of the gallbladder and its stones, GBC risk is considerably reduced.^6^

Epidemiological studies have shown that ethnicity plays a major role in the prevalence of GSD, which is low in Asian and African countries (<10%), higher in Europe (~20%) and reaches >40% in America.^2^ Notably, prevalence in adult Chilean Amerindians is estimated to be 49.4% and 12.3% in native Mapuche women and men, respectively.^7^ Likewise, 36.7% and 13.1% of adult Chilean women and men, admixed from Europeans (mainly from Spain) and indigenous Amerindians (mainly Mapuche Native Americans), are at risk of developing the disease.^7^ Studies in families and case-controls populations have shown that GSD has a strong genetic component.^8 9^ However GWAS and meta-analyses have reported only a handful of lithogenic variants,^10^ the most important within the hepatic sterolin transporter *ABCG8* gene (p.D19H; rs11887534),^11^ which produces a gain-of-function transporter responsible for hyper secretion of cholesterol-saturated bile.^12^ This signal has been replicated in different populations across the world, including admixed Chileans,^11 13 14^however it only explains a small portion of the population attributable risk (PAR=11.2%) of the disease.^15^

Here we report the results of the first large GWAS for a prevalent complex disease in the admixed Chilean population. We hypothesized that a high-density GWAS of GSD in admixed Latinos could identify population specific variants, define the GSD genetic landscape and reveal novel pathological mechanisms.

## METHODS

### Study participants

#### Discovery stage

The Chilean GWAS discovery cohort consisted in 1,235 Chilean individuals, belonging to the ANCORA family health centers in Santiago de Chile (Puente Alto, La Florida and La Pintana). This sample constitutes an admixed (European-Amerindian) group aged between 20-80 years old, of middle-low income, from an urban area and representative of the general Chilean population. Disease status was determined by abdominal ultrasonography for all individuals. Cases were defined as patients with reported cholecystectomy or by the presence of gallstones in their gallbladder. Control individuals had normal gallbladders, neither a history of gallbladder surgery nor gallstone findings in ultrasound. Since women have a higher risk to develop gallstones compared to men, the cohort was mainly composed of women (1,013 individuals: 489 cases and 524 controls), corresponding to the 93.05% of the sample. After genotyping and QCs (see ahead), 1,095 individuals remained (529 cases and 566 controls) in this GWAS discovery cohort (table 1). To reduce variability in the genetic association analyses, cases and controls were matched for known GSD covariates and comorbidities, including body mass index (BMI, 29.44±4.28 and 29.66±3.94, respectively), age (51.32±10.67 and 49.87±9.57, respectively) and sex (92.5% of women), and tested for statistical deviation in cases compared to controls. We excluded Type-2 Diabetes (T2D) individuals from the Discovery stage in order to diminish the presence of confounding factors.

**Table 1.** Clinical characteristics of patients in the Discovery stage and independent replication cohorts

| Variable | Discovery<sup>a</sup> |  | Replication<sup>a</sup> |  | Secondary replication cohorts<sup>b</sup> |  |  |  |  |
| --- | --- | --- | --- | --- | --- | --- | --- | --- | --- |
|  | Stage 1 (n=1,095) |  | Stage 2 (n=1,643) |  | Chile GBC | Germany POPGEN-KIEL (n=1,938) |  | Germany SHIP (n=4,154) |  |
|  | Cases (n=529) | Controls (n=566) | Cases (n=626) | Controls (n=1,017) | Cases (n=397) | Cases (n=1,027) | Controls (n=911) | Cases (n=882) | Controls (n=3,272) |
| Age (years)<sup>c</sup> | 51.3±10.7 | 49.9±9.6 | 59.6±12.5 | 48.3±12.2 | 62.7±12.9 | 45.5±12.3 | 59.4±12.8 | 61.0±13.1 | 47.1±15.9 |
| Sex (% females) | 92.4 | 92.6 | 84.8 | 63.1 | 100 | 60.6 | 42.0 | 65.5 | 47.0 |
| BMI<sup>c</sup> | 29.4±4.3 | 29.7±3.9 | 30.7±5.9 | 28.6±5.4 | N.A. | 27.6±5.3 | 26.5±4.1 | 28.0±3.9 | 26.7±4.5 |
| T2D (% affected) | 0 | 0 | 40.7 | 14.4 | N.A. | N.A. | N.A. | 17.4 | 6.4 |

<sup>a</sup> Chilean Discovery and Replication cohorts are described in Methods.
<sup>b</sup> Secondary replication cohorts correspond to analyses to measure TRAF3 association in different populations where not all necessary data (covariates) were available for adjustment.
<sup>c</sup> All quantitative measures are shown as average and its standard deviation; T2D, Type 2 diabetes; N.A., not available.

#### Chilean GSD replication stage

The independent replication cohort consisted of 1,643 Chilean individuals (626 cases and 1,017 controls), also coming from ANCORA family health centers or a population base study in La Florida, which has been previously described.^7^ As described above, information regarding age, sex, BMI and T2D was gathered for all individuals to control for covariates/comorbidities in the genetic association analyses.

#### Chilean gallbladder cancer (GBC) cohort

A secondary Chilean replication cohort was derived from GBC patients, who almost all (90%) had gallstones. The cohort is composed of a total number of 397 incident and prevalent women cases, recruited from Hospital Sótero del Río, Clinical Hospital of the P. Catholic University, Hospital Regional de Valdivia, Hospital Regional de Temuco and Hospital Regional de Concepción. DNA samples were obtained for incident cases (alive patients) from peripheral blood using the Invisorb^®^ Blood Universal Kit (Stratec, Berlin, Germany). For prevalent cases, DNA was extracted from normal (non-tumoral) tissue samples using the QIAamp DNA-Mini-Kit © (Qiagen, Mainz, Germany). GBC phenotype was defined according to criteria and steps described by the Classification of Malignant Tumors and the American Joint Committee on Cancer (TNM-AJCC) where only adenocarcinomas were included. Each histological cut was reviewed and selected by two independent expert pathologists. After delimiting the healthy tissue, either adjacent to the tumor or other non-tumoral tissues such as liver or cystic ganglion, blocks were selected for the laminae or sequential cuts of 10pm thickness were performed for dissection for extraction of genomic DNA. DNA concentration and integrity were determined by the NanoDrop ND-1000^®^ Spectrophotometer and by agarose gel electrophoresis, respectively.

#### European replication cohorts

We analyzed two European GSD cohorts from Germany. The first cohort comes from the POPGEN biobank of the Medicine Faculty at the University of Kiel, consisting in 1,938 individuals (1,027 cases and 911 healthy controls), genotyped with the Affymetrix Genome-wide Axiom CEU microarray.^16^ Considering that GSD prevalence increases with age,^11^ the statistical analysis performed on this cohort did not include age as a covariate, since stone-free controls were chosen and they had a higher median age (59.43 years old) than affected individuals (45.5 years old). The second cohort comes from the Study of Health in Pomerania (SHIP) from the Greifswald region in northeast Germany, comprising 4,154 subjects (882 GSD cases and 3,272 healthy controls). Genome-wide genotyping was performed with the Affymetrix 6.0 microarrays.^17^ Both cohorts were similarly screened for the presence or absence of GSD using abdominal ultrasonography.

### Genotyping and QC Procedures in the Discovery GWAS Dataset

Main experimental procedures involved in the present study are summarized in figure S1. Genomic DNA was extracted from peripheral blood using (Invisorb Blood Universal kit, *Invitek,* Berlin, Germany). Extracted DNA was quantified with a nanodrop spectrophotometer to a fixed concentration of 50 ng/μ1. Genome-wide genotyping on the discovery cohort was performed with the Affymetrix Axiom^®^ Genome-Wide LAT 1 World Array 4at the Cologne Center for Genomics (University of Cologne, Germany), since the microarray includes 818,155 SNPs and it has been optimized to enhance the capture of variants in populations with Amerindian ancestry.^18^ Genotype intensity files (CEL files) were converted into actual genotypes following the Affymetrix Power tools protocol. Briefly, 1,235 CEL files (1 per sample) containing 818,155 genotypes were processed with the Affymetrix Power Tools and SNPolisher softwares v. 1.17.0 (https://www.thermofisher.com/cl/es/home/life-science/microarray-analysis/microarray-analysis-partners-programs/affymetrix-developers-network/affymetrix-power-tools.html).

### Global Ancestry Estimation

Genetic admixture contributes to the susceptibility of complex diseases.^19^ Since the Chilean population is a mixture of mainly Spanish and Mapuche Native American ancestries, we calculated global ancestry proportions for each individual in the discovery cohort with ADMIXTURE v. 1.3.0 (https://www.genetics.ucla.edu/software/admixture/download.html using the reference panel of the 1000 Genomes project Phase-3 and 11 individuals from the Mapuche-Huilliche ethnic group as a Chilean Native American reference panel (HUI).^20^ The mean proportion of the Amerindian ancestry and its standard deviation, calculated in cases and controls, was tested for significant deviation using a one-way ANOVA test.

### Functional characterization of TRAF3

We performed gene expression analysis for TRAF3 at mRNA and Protein level in gallbladder and duodenal mucosa tissue samples coming from GSD and control individuals. We also carried out histological and immunohistochemical analysis of the TRAF3 protein in gallbladder tissue samples.

Additional information with details of the presented methods, including GWAS QCs, genotvpe imputation, replication, statistical analyses and functional gene expression studies is provided in the Supplementary Data.

## RESULTS

### Discovery GWAS for GSD in an admixed Chilean population

The Discovery GWAS stage involved 1,235 admixed Chileans Latinos with Mapuche Native American Ancestry genotyped using the Affymetrix Axiom^®^ Genome-Wide LAT 1 World Array 4 (figure S1). After genotype calling and quality controls, 1,095 individuals (529 GSD cases and 566 controls) and 677,835 SNPs remained for analysis (table 1). We imputed genotypes in these individuals using the 1000 Genomes Project Phase 3 reference panel and achieved a total number of 9,433,911 SNPs and Indels. We tested for genetic association with GSD by performing a logistic regression analysis adjusted by age (cases=51.32±10.67, controls=49.87±9.57, one-way ANOVA P=0.018), since other known associated covariates such as sex (cases=92.43%, controls=92.57%, Z score test P=0.928), BMI (cases=29.44±4.28, controls=29.66±3.94, one-way ANOVA P=0.376) and ancestry composition (cases=46.3%±7.0%, controls=45.5%±7.5%, one-way ANOVA P=0.069), did not differ significantly between cases and controls.

Genome-wide association results are presented in the figure 1 and consider a genomic control inflation factor λ=1.02 (figure S2). While we did not detect genome-wide significant associations, several suggestive signals (P<1×10^−5^) were observed. Top-ten candidate variants with suggestive association with GSD were found within *ELMO1* (rs4446645, P=3.36×10^−7^), 5q34 locus (rs10463138, P=2.47×10^−6^), 4q12 locus (rs74537816, P=3.54×10^−6^), *TRAF3* (rs368550004, P=3.92×10^−6^), 9p21.1 locus (rs4879592, P=4.66×10^−6^), *OLFML2B* (rs10918361, P=4.91×10^−6^), 3p22.2 locus (rs73827633, P=5.06×10^−6^), *ABCG8* (rs11887534, P=5.24×10^−6^), 11p15.3 locus (rs147367002, P=7.63×10^−6^) and *TRPV1* (rs7223530, P=8.07×10^−6^) (table 2). Aside from *ABCG8,* none of the GWAS signals associated with GSD reported in a recent large meta-analysis of individuals with European ancestry^10^ reached suggestive genome-wide significance (table S1).

**Table 2.** Top-10 candidate variants associated with GSD in admixed Chileans in the Discovery and Replication stages

| SNP | Locus | Chr | SNP ID | RA | RAF | Discovery |  | Replication |  | Combined |  |  |
| --- | --- | --- | --- | --- | --- | --- | --- | --- | --- | --- | --- | --- |
|  |  |  |  |  |  | P value | OR (95% CI) | P value | OR (95% CI) | MP value<sup>b</sup> | I<sup>2c</sup> | OR (95% CI) |
| SNP 1 | *ELMO1* | 7 | rs4446645 | A | 0.66 | $3.4 \times 10^{-7}$ | 1.6 (1.3-1.8) | 0.72 | 1.0 (0.9-1.2) | - | - | - |
| SNP 2 | 5q34 | 5 | rs10463138 | A | 0.52 | $2.5 \times 10^{-6}$ | 1.5 (1.3-1.8) | 0.43 | 0.9 (0.8-1.1) | - | - | - |
| SNP 3 | 4q12<sup>d</sup> | 4 | rs74537816 | T | 0.95 | $3.5 \times 10^{-6}$ | 2.3 (1.6-3.3) | - | - | - | - | - |
| | | 4 | rs1824387 | A | 0.96 | $4.0 \times 10^{-5}$ | 2.1 (1.5-2.9) | 0.55 | 1.1 (0.8-1.5) | - | - | - |
| SNP 4 | *TRAF3*<sup>e</sup> | 14 | rs368550004 | G | 0.69 | $3.9 \times 10^{-6}$ | 1.5 (1.3-1.8) | 0.009 | 1.3 (1.1-1.5) | - | - | - |
| | | 14 | rs12882491 | C | 0.69 | $5.3 \times 10^{-6}$ | 1.5 (1.3-1.8) | 0.003 | 1.3 (1.1-1.5) | $1.1 \times 10^{-7}$ | 24.7 | 1.4 (1.2-1.6) |
| SNP 5 | 9p21.1 | 9 | rs4879592 | C | 0.86 | $4.7 \times 10^{-6}$ | 1.7 (1.3-2.1) | 0.35 | 1.1 (0.9-1.4) | - | - | - |
| SNP 6 | *OLFML2B* | 1 | rs10918361 | G | 0.43 | $4.9 \times 10^{-6}$ | 1.5 (1.3-1.8) | 0.09 | 0.9 (0.7-1.0) | - | - | - |
| SNP 7 | 3p22.2 | 3 | rs73827633 | T | 0.98 | $5.1 \times 10^{-6}$ | 3.0 (1.9-4.8) | 0.73 | 0.9 (0.6-1.4) | - | - | - |
| SNP 8 | *ABCG8* | 2 | rs11887534 | C | 0.14 | $5.2 \times 10^{-6}$ | 1.9 (1.4-2.5) | 0.001 | 1.6 (1.2-2.1) | $3.2 \times 10^{-8}$ | 0 | 1.7 (1.5-2.0) |
| SNP 9 | 11p15.3<sup>d</sup> | 11 | rs147367002 | A | 0.14 | $7.6 \times 10^{-6}$ | 1.9 (1.5-2.6) | - | - | - | - | - |
| | | 11 | rs16908929 | A | 0.15 | $4.3 \times 10^{-5}$ | 1.8 (1.3-2.3) | 0.47 | 1.1 (0.9-1.4) | - | - | - |
| SNP 10 | *TRPV1* | 17 | rs7223530 | G | 0.77 | $8.1 \times 10^{-6}$ | 1.6 (1.3-1.9) | 0.27 | 0.9 (0.7-1.1) | - | - | - |

<sup>a</sup>Combined analysis correspond only to the variants surpassing Bonferroni correction ( $P < 0.005$ ) at replication stage.
<sup>b</sup>MP value (Meta-analysis P value) correspond to Fixed effect estimates.
<sup>c</sup>I<sup>2</sup> indicates between-cohort heterogeneity.
<sup>d</sup>The leading variants selected for the candidates weren't genotyped by TaqMan in the replication cohort due technical issues, and were replaced by adjacent SNPs in high LD: rs1824387 for SNP 3, $r^2=0.88$ ; and rs16908929, $r^2=0.88$ ; respectively.
<sup>e</sup>For *TRAF3* we genotyped the leading variant rs368550004 (Indel) which is the best SNP in high LD rs12882491, $r^2=1.0$ . RA: risk allele; RAF: risk allele frequency; OR: odds ratio; CI: confidence interval.

**Figure 1.**
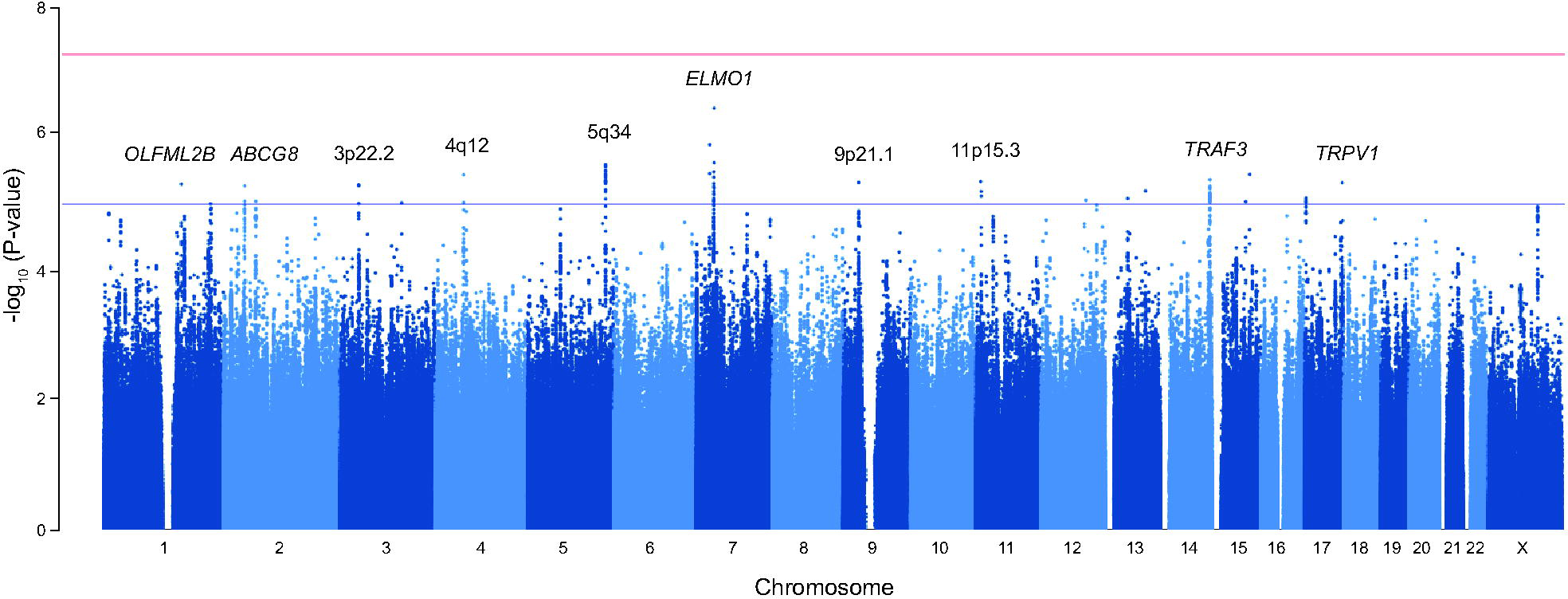
Genome-wide association results for GSD in admixed Chilean in the Discovery stage. Manhattan plot depicting the association P values for all good quality variants. Red line shows genome-wide significance threshold (P<5×10^−8^). Top-ten candidate variants surpassing the suggestive genome-wide significance threshold (blue line, P<1×10^−5^) are shown, which were taken for further replication.

### Association of *ABCG8* and *TRAF3* variants with GSD in an independent Chilean cohort

The top-ten candidate variants with suggestive association with GSD in the Discovery stage were tested for replication in an independent admixed Chilean cohort composed of 1,643 individuals (626 cases and 1,017 controls) from Santiago de Chile. Replication was performed by real-time qPCR with TaqMan probes for most variants, with the exception of SNP3 (4q12), SNP4 *(TRAF3)* and SNP9 (11p15.3), which were examined by proxy SNPs in high linkage disequilibrium (LD; r^2^>0.88). After genotyping and logistic regression analyses adjusted for age (cases=59.57±12.47, controls=48.26±12.19, one-way ANOVA P<0.001), sex (cases=84.82%, controls=63.12%, Z score test P<0.001), BMI (cases=30.74±5.92, controls=28.61±5.38, one-way ANOVA P<0.001) and T2D (cases=40.73%, controls=14.36%, Z score test P<0.001), we replicated variants within the *ABCG8* (rs11887534, P=0.001) and *TRAF3* (rs12882491, P=0.003) genes (figure 2, table 2). Both variants passed Bonferroni correction for 10 tests (P<0.005) and displayed an effect with the same direction as it was observed in the Discovery stage of the GWAS.

**Figure 2.**
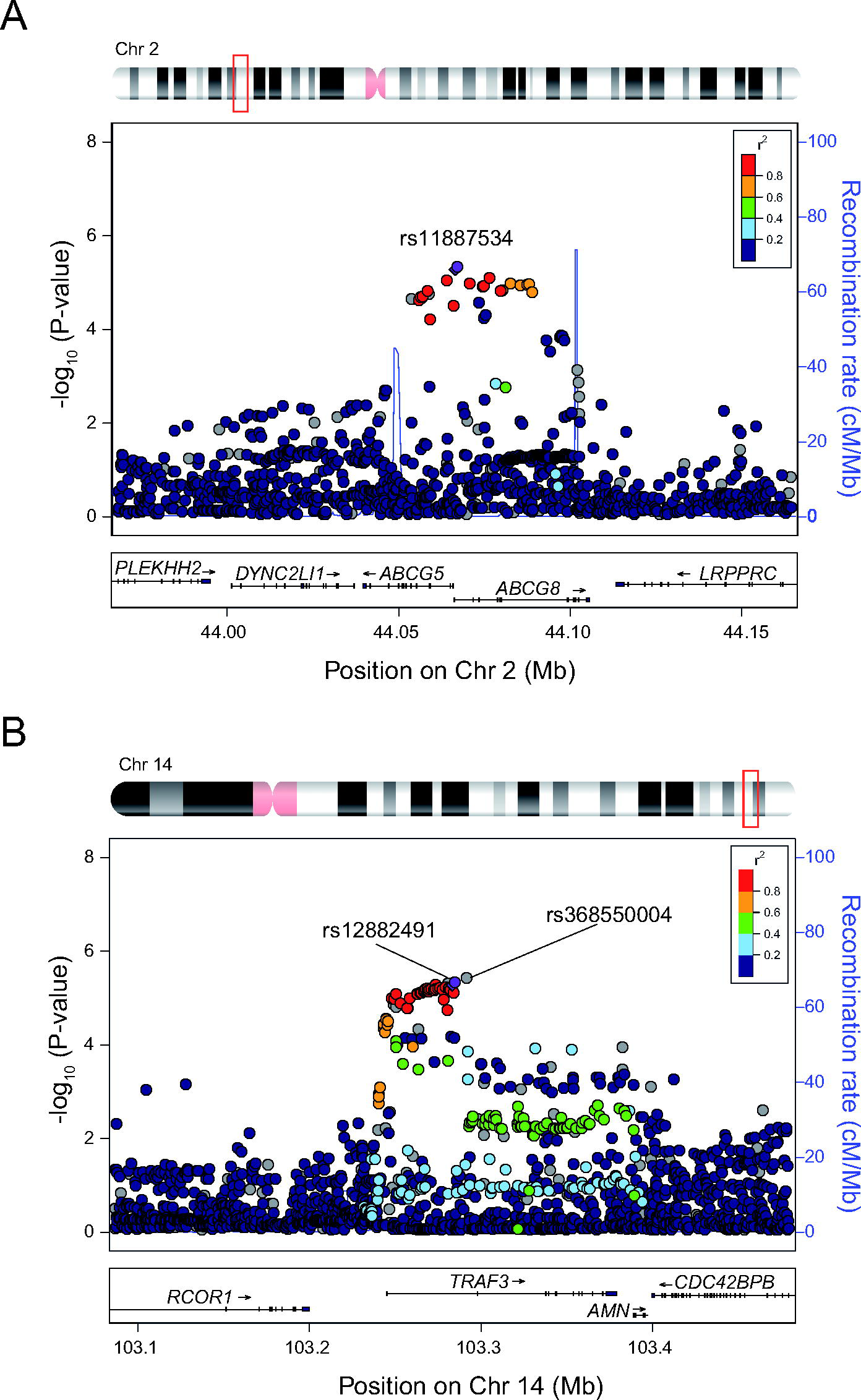
Regional association plots for the *ABCG8* and *TRAF3* association signals in the GWAS Discovery stage. Locus zoom for *ABCG8* (A) and *TRAF3* (B) signals in chromosome 2 and 14, respectively. To the left is the P value related to GSD in a log10 scale. SNPs with the highest association to GSD are colored purple. The insert panel denotes the imputed quality score (r2) and the appropriate SNPs are colored in the plot.

To assess for the combined effect between *ABCG8* and *TRAF3* SNPs with GSD we performed meta-analysis^22^ in the Discovery and Replication stages and observed that association with GSD increased for *ABCG8* (rs11887534, P=3.24×10^−8^, OR=1.74) and *TRAF3* (rs12882491, P=1.11×10^−7^, OR=1.40) variants (table 2). These results further confirm *ABCG8* as a risk factor for GSD and identify *TRAF3* as a novel gene associated with the disease in the Chilean Admixed population.

### *ABCG8* and *TRAF3* variants are associated with GBC

Although GBC is the most severe complication of GSD,^5 6 23^ few studies have investigated the effect of *ABCG8* as a genetic determinant for the disease.^24 25^ We therefore examined the orientation and magnitude of the effect for *ABCG8* and *TRAF3* variants in a Chilean cohort of GBC patients (n=397 women), using sex-matched controls from the Replication population (n=667 women). After logistic regression analyses adjusted by age (cases=62.65±13.08, controls=43.38±12.13, one-way ANOVA P<0.001), we observed that both SNPs were associated with GBC (*ABCG8* rs11887534: P=6.9×10^−4^, OR=1.77; *TRAF3* rs12882491: P=0.045, OR=1.24) (table 3), with same direction of the effect and similar risk allele frequency (RAF), as it was observed in the Discovery and Replication cohorts (RAF_*ABCG8*_: GBC=0.14, Discovery=0.14, Replication=0.12; RAF_*TRAF3*_: GBC=0.66, Discovery=0.69, Replication=0.66) (figure 3). These results confirm the genetic association of *ABCG8* with GBC and reveal *TRAF3* as a novel marker for the disease.

**Figure 3.**
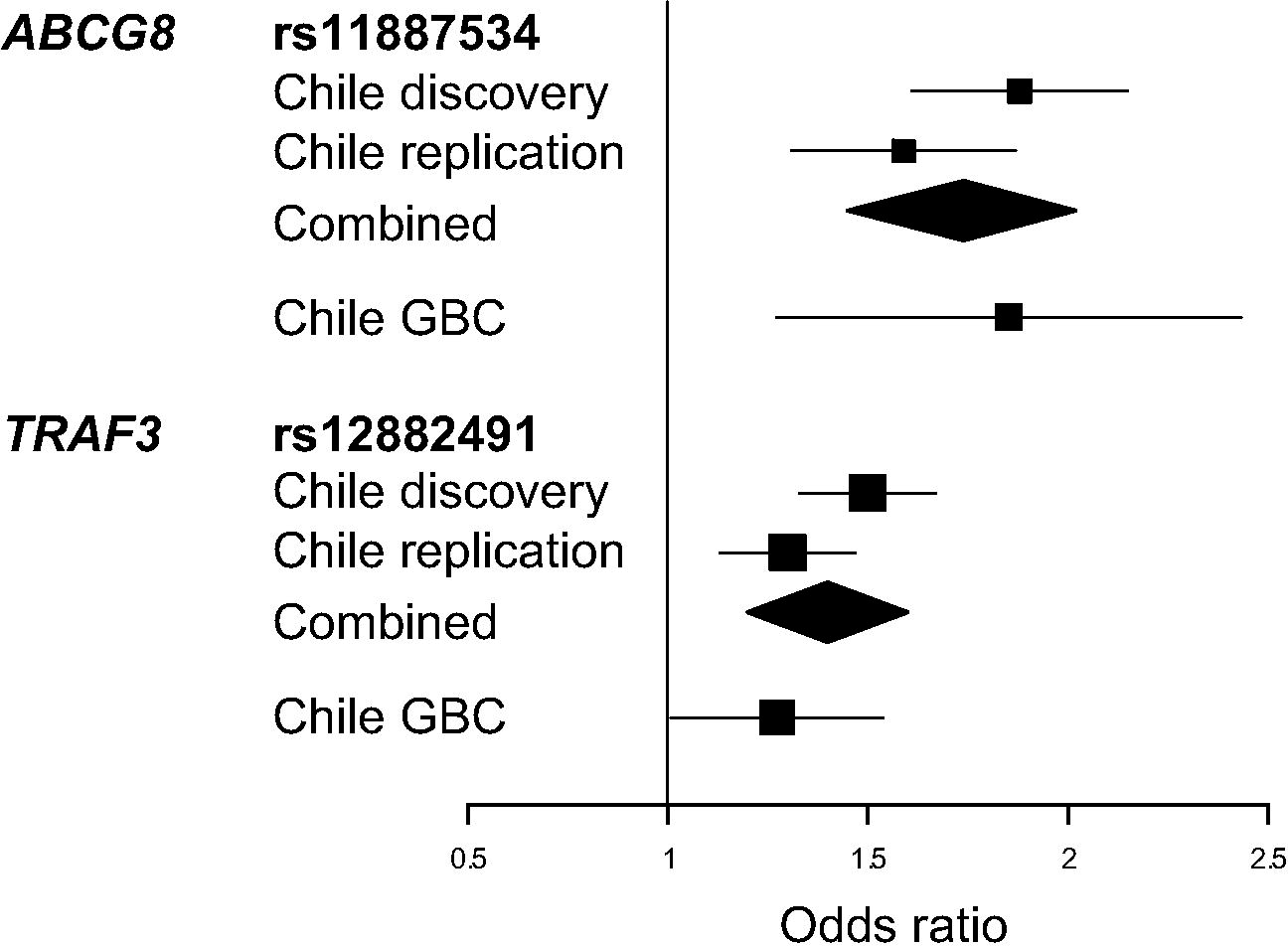
Effect size for *ABCG8* and *TRAF3* association signals in admixed Chilean replication cohorts. Forest plot calculated for GSD and GBC cohorts. The effect is shown in OR values and their 95% confidence intervals.

**Table 3.** *ABCG8* and *TRAF3* variants are associated with GBC

| Gene | SNP | RAF | P value | OR (CI 95%) |
| --- | --- | --- | --- | --- |
| *ABCG8* | rs11887534 | 0.14 | $6.9 \times 10^{-4}$ | 1.8 (1.3-2.5) |
| *TRAF3* | rs12882491 | 0.66 | 0.045 | 1.2 (1.0-1.5) |

RAF = Risk allele frequency; OR: odds ratio; CI: confidence interval.

**RAF = Risk allele frequency; OR: odds ratio; CI: confidence interval.**

### Replication Analyses for *ABCG8* and *TRAF3* variants in Europeans cohorts

We originally identified *ABCG8* as a risk factor for GSD in a German population from Kiel.^11 12^ Therefore, we examined *ABCG8* rs11887534 and *TRAF3* rs12882491 variants in an extended sample from the original POPGEN-Kiel cohort consisting in 1,938 individuals (1,027 cases and 911 controls),^16^ as well as in 4,154 individuals (882 cases and 3,272 controls) from the north-east side of Germany identified as the SHIP-Greifswald^17^ cohort (table 1). We found that *ABCG8* rs11887534 replicated in both POPGEN-Kiel (P=5.36×10^− 10^, OR=2.00) and SHIP-Greifswald (P=1.29×10^−6^, OR=1.87) cohorts. Conversely, no replication was observed for *TRAF3* rs12882491 in either the POPGEN-Kiel (P=0.65, OR=0.97) or the SHIP-Greifswald (P=0.39, OR=1.06) cohorts (table 4). A difference in orientation and/or magnitude of the effect with the Chilean cohorts was also reflected in terms of risk allele frequency (RAF_*TrAF3*_ POPGEN-Kiel=0.34, SHIP-Greifswald=0.35). Altogether, these results demonstrate a consistent effect of *ABCG8* association with GSD in different world populations and indicate an ethnic-specific effect for the *TRAF3* variant in admixed Latinos with Native Amerindian ancestry.

**Table 4.** Association Results for *ABCG8* and *TRAF3* SNPs in the German GSD cohorts

| Population | ABCG8 rs11887534 |  |  | TRAF3 rs12882491 |  |  |
| --- | --- | --- | --- | --- | --- | --- |
|  | RAF | P value | OR (CI 95%) | RAF | P value | OR (CI 95%) |
| POPGENE-Kiel | 0.12 | $5.4 \times 10^{-10}$ | 2.0 (1.6-2.5) | 0.34 | 0.65 | 1.0 (0.8-1.1) |
| SHIP-Greifswald | 0.08 | $1.3 \times 10^{-6}$ | 1.9 (1.5-2.4) | 0.35 | 0.39 | 1.1 (0.9-1.2) |

RAF = Risk allele frequency; OR: odds ratio; CI: confidence interval

RAF = Risk allele frequency; OR: odds ratio; CI: confidence interval

### TRAF3 expression in human gallbladder and duodenal samples

*TRAF3* is a member of the tumor necrosis factor (TNF) receptor-associated factor (TRAF) protein family that mediates signal transduction during the activation of immune and inflammatory responses.^26-28^ To date, seven *TRAF* genes are known in humans (*TRAFI* to *TRAF7*) and the products encoded by these genes are involved in cytokine production and cell survival.^29^ Considering that acute and chronic inflammation (cholecystitis) are known hallmarks of GSD pathophysiology,^30^ we explored the expression of the *TRAF3* gene in gallbladder and duodenal tissues resected from cases and control individuals. Immunohistochemical staining in the gallbladder mucosa revealed that TRAF3 protein was observed in different gallbladder cell types such as the epithelium, smooth muscle fibers, arterioles and veins (figure 4A). Notably, we found a significant decrease in TRAF3 protein levels in the duodenal mucosa of GSD cases compared to control individuals (P<0.001, two-tailed t-test; figure 4B and 4C) and also a significant decrease of *TRAF3* transcripts in a set of 41 RNA samples from duodenal (19 cases, 22 controls, P<0.001, two-tailed t-test) and gallbladder mucosa (11 cases and 11 controls, P=0.015, two-tailed t-test) (figure 4D). These results show significant lower levels of TRAF3 in GSD affected individuals, suggesting a functional contribution of this gene during the inflammatory response observed during the onset or development of the disease.

**Figure 4.**
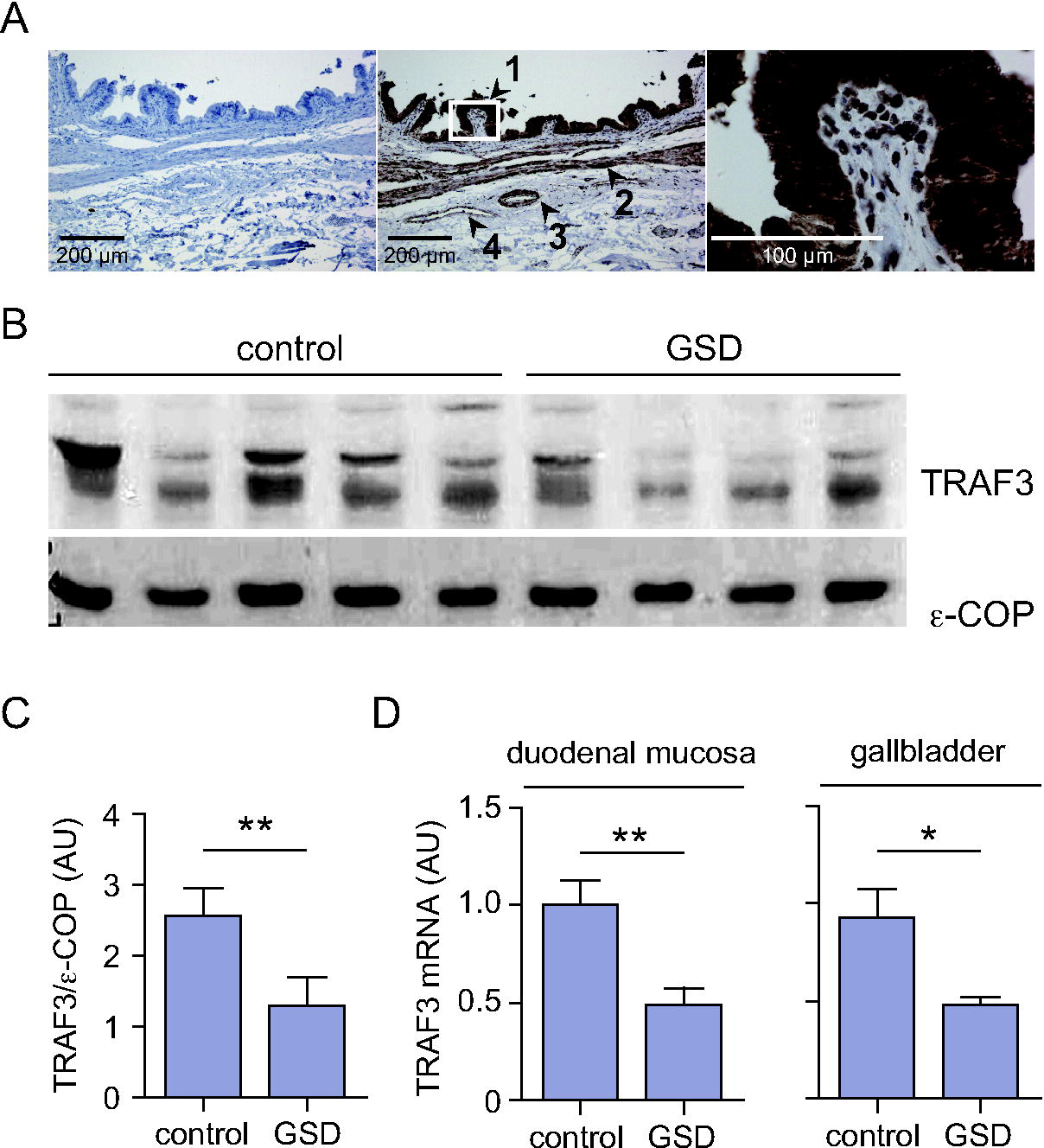
*TRAF3* expression in human gallbladder and duodenal tissues. (A) Immunohistochemical analysis of TRAF3 expression in human gallbladder. Left panel: Negative control without primary antibody; Middle panel: Positive staining was observed for TRAF3 in the cell epithelium (1), muscle fibers (2), arterioles (3), and veins (4); Right panel: Zoom-in (60x) showing TRAF3 localization in the mucosal epithelium. (B) Western blot for *TRAF3* in biopsies of duodenal mucosa tissue from GSD cases and control individuals. (C) Determination of TRAF3 protein levels in the duodenal mucosa (as is observed in B). (D). Differential expression analyses for TRAF3 in duodenal (left panel) and gallbladder mucosa tissues (right panel) in GSD cases and control samples.

## DISCUSSION

Herein we report that common genetic variation in *ABCG8* and *TRAF3* genes confer risk to GSD and GBC in the admixed Chilean population. The signal replicated for the *ABCG8* gene has been reported in several world populations including Latinos.^10^ In the Discovery stage, the leading SNP rs11887534 (D19H) showed a lower magnitude of association, compared with the estimated from the German population,^11^ and this could be explained by the difference in the risk allele frequency (C allele) that is higher in Chilean controls (0.079 vs. 0.05, respectively). This difference is also reflected in the effect size, where a lower OR in Chile (OR = 1.88, CI 95% = 1.43-2.47) is observed compared with Germany (OR = 2.00, CI 95% 1.61-2.50). This could be a reason for not being able to detect in the discovery stage a genome-wide signal (p < 5×10^−8^) in the *ABCG8* locus, suggesting that larger sample sizes are required to reach that significance level in future studies on this population.

The OR obtained for *ABCG8* in the discovery stage represents a PAR of 8.7%, while for the *TRAF3* signal (rs12882491, risk allele C) the OR of 1.5 (CI 95% = 1.26-1.78) represents a PAR of 24.2%, indicating that *TRAF3* has a higher genetic contribution than *ABCG8* in this cohort. Taking together, the combined effect of both associations gives an estimated PAR of 30.79%, meaning that if their contribution for the disease is hypothetically eliminated^31^, there would be 30.79% fewer affected individuals in the population. In addition, SNP x SNP epistasis analysis showed non-significant interaction between *ABCG8* and *TRAF3* association signals (p = 0.399), suggesting that the genetic contribution of both risk factors is independent from each other. While the association *ABCG8* variant was consistently observed in all the populations in this study, *TRAF3* association is only present in Chilean samples, indicating an ethnic-specific effect. This needs to be further validated in other closely related Latino populations from Peru, Argentina and Bolivia that also have a high GSD prevalence.^32-34^

Here we also observed that *ABCG8* and *TRAF3* variants are significantly associated with GBC in admixed Chileans. The *ABCG8* rs11887534 variant has been previously associated with this malignancy in China^25^, India^24^ and European populations^35^ and most of the studies show that the GBC risk is more pronounced in patients that also had biliary stones in the gallbladder, as it is likely observed in this Chilean cohort (i.e. 90% of female GBC patients with cholesterol gallstones). This is the first time that the *ABCG8* and *TRAF3* genes showed significant association with GBC in an American population and thus *TRAF3* represents a novel risk factor for this malignancy.

TRAF3 belongs to the TNF receptor associated factor (TRAF) protein. TRAF3 function has been related to the promotion of autoimmunity and predisposition to various cancers,^36^ including multiple myeloma,^37^ Hodgkin lymphoma^38^ and splenic neoplasms.^39^ The protein physically interacts with the NF-κB-inducing kinase (NIK) promoting NIK degradation by the proteasome, which results in the inactivation of the non-canonical NF-κB pathway.^40^ Indeed, siRNA knockdown of *TRAF3* in human cancer cell lines stabilizes NIK and activates NF-κB dependent transcription.^41^ Interestingly, recent evidence indicate that activation of NF-κB results in invasion, lymphangiogenesis and tumor growth in GBC tissue and cell lines,^42 43^ and therefore genetic alterations in one of its modulators could be directly involved in GBC pathogenesis and may serve as a target of future therapies and prevention.

Inflammation is a hallmark of GSD pathogenesis. Maurer et al reviewed the roles of infection, inflammation and the immune system in cholesterol gallstone formation and suggested that the formation of biliary stones is preceded by histopathologic alterations in the gallbladder wall that indicate inflammation,^44^ including edema, increased wall thickness, decreased motility and altered transport function in the epithelium. That evidence indicates a crosstalk between the formation of cholesterol stones and inflammatory processes that promote the development and progression of the disease and suggests that *ABCG8* and *TRAF3* could be both relevant actors in GSD pathology. First, it is known that the *ABCG8* variant promotes gallstone formation by enhancing the efflux of cholesterol-saturated bile.^12^ Second, lower expression levels of TRAF3 in duodenal mucosa and gallbladder epithelium of GSD patients (as described in the present study) could be responsible for increased production of pro-inflammatory factors, such as NF-κB, IL-6 and IL-12B, as well as for decreased activity of anti-inflammatory factors such as type 1 interferon and IL-10.^41 45-47^ The latter has already been associated with GSD and GBC development and progression.^48^ In addition, a *TRAF3* (R118W) mutation that correlates with lower expression levels of the gene, has been associated with higher susceptibility to the development of multiple myelomas and herpes simplex virus encephalitis,^49 50^ further stressing that variants controlling TRAF3 mRNA levels may be involved in GSD and GBC etiology.

In summary, our results confirm the association of *ABCG8* and identify *TRAF3* as a novel candidate gene for GSD and GBC in the high-risk Chilean population. These genes could be functionally involved in the cascade of events that triggers gallstone formation and the inflammatory response observed in affected patients. Our results may help to develop novel therapies and/or prevention strategies for these prevalent diseases.

## ACKNOWLEDGEMENTS

We gratefully acknowledge affected and unaffected individuals who participated in this study.

## AUTHOR CONTRIBUTIONS

B.I.B., G.V.D and J.F.M. conceived and designed the experiments. J.F.M., W.L., A.F., S.H., G.B., W.v.S., C.S., H.V., U.V., G.H., M.M.L., K.P. and C.B., collected the data. B.I.B., E.P.-P., L.A., S.B., M.T., E.R., G.D.U and G.V.D. performed experiments or analyzed the data. P.N., J.L.S., R.A.G. and J.H. helped in drafting the manuscript. B.I.B., G.V.D and J.F.M. wrote the paper. All authors reviewed the manuscript for medical and scientific accuracy.

## DATA AVAILABILITY

All genotype files for the Chilean Discovery population are be available upon request to the corresponding authors.

## ETHICAL STATEMENT

All procedures involving genetic and clinical data usage from Chilean GSD and GBC patients were approved by the Ethical Scientific Committee at the Pontificia Universidad Catolica de Chile and were conducted in accordance with the guidelines of the National Commission on Science and Technology (CONICYT-Chile). All participants provided informed consent. In the German replication cohorts, Ethical Scientific Committees have previously approved these studies ^16 17^

